# Nanobodies identify an activated state of the TRIB2 pseudokinase

**DOI:** 10.1101/2022.04.29.489987

**Authors:** Sam A Jamieson, Michael Pudjihartono, Christopher R Horne, Robert C Day, James M Murphy, Peter D Mace

## Abstract

Tribbles proteins (TRIB1–3) are a pseudokinase-only branch of the human kinome, which recruit substrates to the COP1 ubiquitin-ligase for ubiquitination. TRIB2 was the first Tribbles ortholog to be implicated as a myeloid leukaemia oncogene, by way of recruiting the C/EBPa transcription factor for degradation by COP1. Here we report selection and characterisation of nanobodies against the TRIB2 pseudokinase domain from a synthetic yeast surface-display library. We identified nanobodies that bind the TRIB2 pseudokinase domain with low nanomolar affinity. A crystal structure of Nb4.103 in complex with TRIB2 identified a mode of binding to the N-terminal lobe of the pseudokinase, in a manner that enables specific recognition of TRIB2 over TRIB1 and TRIB3. In the nanobody-stabilised state, TRIB2 adopts an activated conformation that is remarkably similar to the C/EBPa-bound state of TRIB1. Characterization in solution revealed that Nb4.103 can stabilise a TRIB2 pseudokinase domain dimer in a face-to-face manner. Conversely, a distinct nanobody (Nb4.101) binds through a similar epitope but does not readily promote dimerization. In combination, this study identifies features of TRIB2 that could be exploited for the development of inhibitors, and nanobody tools for future investigation of TRIB2 function.

## Introduction

Protein kinases are essential mediators of eukaryotic cell biology, with a characteristic fold that binds ATP to mediate substrate phosphorylation (Kornev and Taylor, 2015; Manning et al., 2002). However, approximately 10% of metazoan proteins that contain a kinase domain do not catalyse phosphorylation, and function through diverse alternative mechanisms (Boudeau et al., 2006; Kwon et al., 2019; Mace and Murphy, 2021; Murphy et al., 2014). Most commonly, pseudokinases allosterically regulate other proteins, act as scaffolds to mediate protein complexes, or act as signaling switches (Mace and Murphy, 2021; Murphy et al., 2017). Tribbles proteins (TRIB1–3) are a family of pseudokinases with essential signalling roles in many eukaryotic cells. In mammals, TRIB2 is the most ancestral member of the family, from which other TRIB homologs (TRIB1 and TRIB3) have evolved (Eyers et al., 2017). The Tribbles pseudokinases share a conserved three-domain architecture consisting of an N-terminal extension, a substrate-binding pseudokinase domain, and a C-terminal extension that contains a binding motif for COP1 ubiquitin ligase (Kung and Jura, 2019a; Qi et al., 2006; Uljon et al., 2016; Yoshida et al., 2013). While various functions for Tribbles proteins have been identified, perhaps the best characterised is their role as adaptors for substrate ubiquitination (Kung and Jura, 2016; Murphy et al., 2015). The substrate adaptor role of Tribbles entails the binding of a substrate to the pseudokinase domain, coupled with the recruitment of a COP1 ubiquitin ligase to the C-terminus. This leads to the COP1-facilitated polyubiquitination of the substrate, leading to its proteasomal degradation. Due to their roles in the regulation of transcription factors and metabolic proteins, Tribbles proteins have been variously implicated in a wide range of pathologies, particularly cancer and cardiometabolic disease (Du et al., 2003; Johnston et al., 2019; Keeshan et al., 2006, 2010; Qi et al., 2006; Quiroz-Figueroa et al., 2021).

TRIB1 and TRIB2 have been associated with human acute myeloid leukemia (AML) for some time (Keeshan et al., 2006; Röthlisberger et al., 2007; Rücker et al., 2006), and TRIB3 has also been recently implicated in the disease (Luo et al., 2020). In mouse models, induced retroviral overexpression of TRIB1 and TRIB2 in bone marrow cells leads to the development of AML (Jin et al., 2007; Keeshan et al., 2006, 2010). The role of TRIB1 and TRIB2 as myeloid oncogenes appear to be driven by their ability to induce excessive degradation of C/EBPα via COP1 (Keeshan et al., 2006; Yoshida et al., 2013; Yoshino et al., 2021). In myeloid cells, C/EBPα is a transcription factor that is essential for the process of granulopoiesis (differentiation of myeloid cells), and is itself subject to frequent mutation in AML (Nerlov, 2004; Pulikkan et al., 2017). TRIB-induced degradation of C/EBPα disrupts myeloid differentiation, thus promotes tumorigenesis and the development of AML. The key role of TRIB1 and TRIB2 in AML and other cancers raises their spectre as potential therapeutic targets (Foulkes et al., 2018; McMillan et al., 2021). Like many pseudokinases however, a more complete understanding of their non-catalytic mechanisms will be key to fully realising this potential (Bailey et al., 2015a; Kung and Jura, 2019b; Mace and Murphy, 2021).

Previous structural knowledge of the Tribbles family has been centred on TRIB1— the crystal structure of TRIB1 pseudokinase has been solved in two different states (Jamieson et al., 2018; Murphy et al., 2015). Initially, structural analysis of the TRIB1 pseudokinase domain alone revealed a unique auto-inhibitory conformation whereby the C-terminal COP1-binding motif of TRIB1 binds to a groove formed by the αC-helix within the pseudokinase domain (Murphy et al., 2015). This interaction is mutually exclusive with the C-terminal COP1-binding motif binding to COP1 (Uljon et al., 2016). Later, structural analysis of the TRIB1-C/EBPα complex showed that TRIB1 underwent a significant conformational change upon C/EBPα binding. Upon C/EBPα binding, the activation loop of the pseudokinase domain becomes fully ordered to form the C/EBPα binding site, and the αC-helix rotates from its initial conformation. Specifically, the C-terminal domain is released from its interaction with the pseudokinase domain, thereby releasing TRIB1’s autoinhibition, and enabling COP1 binding (Jamieson et al., 2018). The most closely related protein to TRIB1 that has been structurally characterised is STK40 (Durzynska et al., 2017), which can also bind to COP1 through a C-terminal extension but has a distinctly structured αC-helix that appears unlikely to undergo a similar inactive-active transition. Taken together, it is evident that TRIB1 exhibits conformational dynamics in a similar vein to conventional kinases (Kornev and Taylor, 2015); and to dynamics recently uncovered in other pseudokinase switches such as MLKL in necroptosis and pseudokinase receptor tyrosine kinases (Davies et al., 2020; Garnish et al., 2021; Sheetz et al., 2020). In all of these cases, stabilization of different active sites conformations by small molecules could modulate function (Kung and Jura, 2019b).

To further understand the potential for specifically modulating proteins within the Tribbles family, we sought to structurally characterise the TRIB2 pseudokinase domain. Previous attempts to work with the isolated protein suggested dynamic instability that, combined with the known dynamics of TRIB1, led us to seek reagents that could stabilise TRIB2. To this end, we turned to protein nanobodies, which are derived from camelid species and are distinct from conventional antibodies because they contain only a heavy chain, rather than heavy and light chains (Beghein and Gettemans, 2017; Uchański et al., 2020). This means that the portion of the nanobody that confers specificity is within three complementarity-determining regions (CDRs) in a single compact domain. Specifically, we used a synthetic yeast-display library developed by McMahon et al. as a platform for nanobody discovery (McMahon et al., 2018). By using the yeast-display method, we identified several nanobodies that bind to TRIB2 with low nanomolar affinity and used one of these to solve the structure of TRIB2. The structure reveals a similar architecture to TRIB1, albeit harbouring specific differences surrounding the pseudoactive site that differentiate TRIB1 and TRIB2. Biochemical characterisation of these nanobodies suggests that they may form the basis of bioactive tools to study TRIB2 in cells.

## Results

### Selection of nanobodies against TRIB2

To generate new tools to investigate TRIB2, we used a synthetic yeast-display library to raise nanobodies against the pseudokinase domain (McMahon et al., 2018). Having tested several TRIB2 construct lengths for expression and solution behaviour, we chose to use a construct containing the pseudokinase domain and the C-terminal COP1-binding motif (residues 53–343). Previous studies have shown that constructs containing the TRIB2 pseudokinase and C-terminal extension have similar thermal stability to full-length protein, whereas further truncations have lower stability (Foulkes et al., 2018). Following recombinant expression of TRIB2(53–343) in *E. coli*, we labelled the protein with fluorescent dyes (FITC and Alexafluor647) and performed successive rounds of magnetic and fluorescent cell sorting at progressively more stringent concentrations (see methods). Analysis of the selected library by next generation sequencing allowed us to identify an array of enriched nanobody sequences (Figure 1A; Supplementary Data), a subset of which also exhibited ideal behaviour upon recombinant expression and size-exclusion chromatography. We then assessed the ability of each nanobody to co-elute with the TRIB2 pseudokinase domain upon size-exclusion chromatography, which we took to be a positive indication of high-affinity binding. By itself, TRIB2(53–343) eluted as a relatively broad and heterogenous peak in buffer conditions with near-neutral pH (dotted traces, Figure 1B), whereas three clones caused a marked improvement in the elution profile into a more homogenous peak, and co-eluted with TRIB2 (Figure 1B). Following initial screening steps, we prioritised three of the co-eluting nanobodies (Nb4.101, Nb4.103, and Nb4.105) for detailed characterisation.

**Figure 1-.**
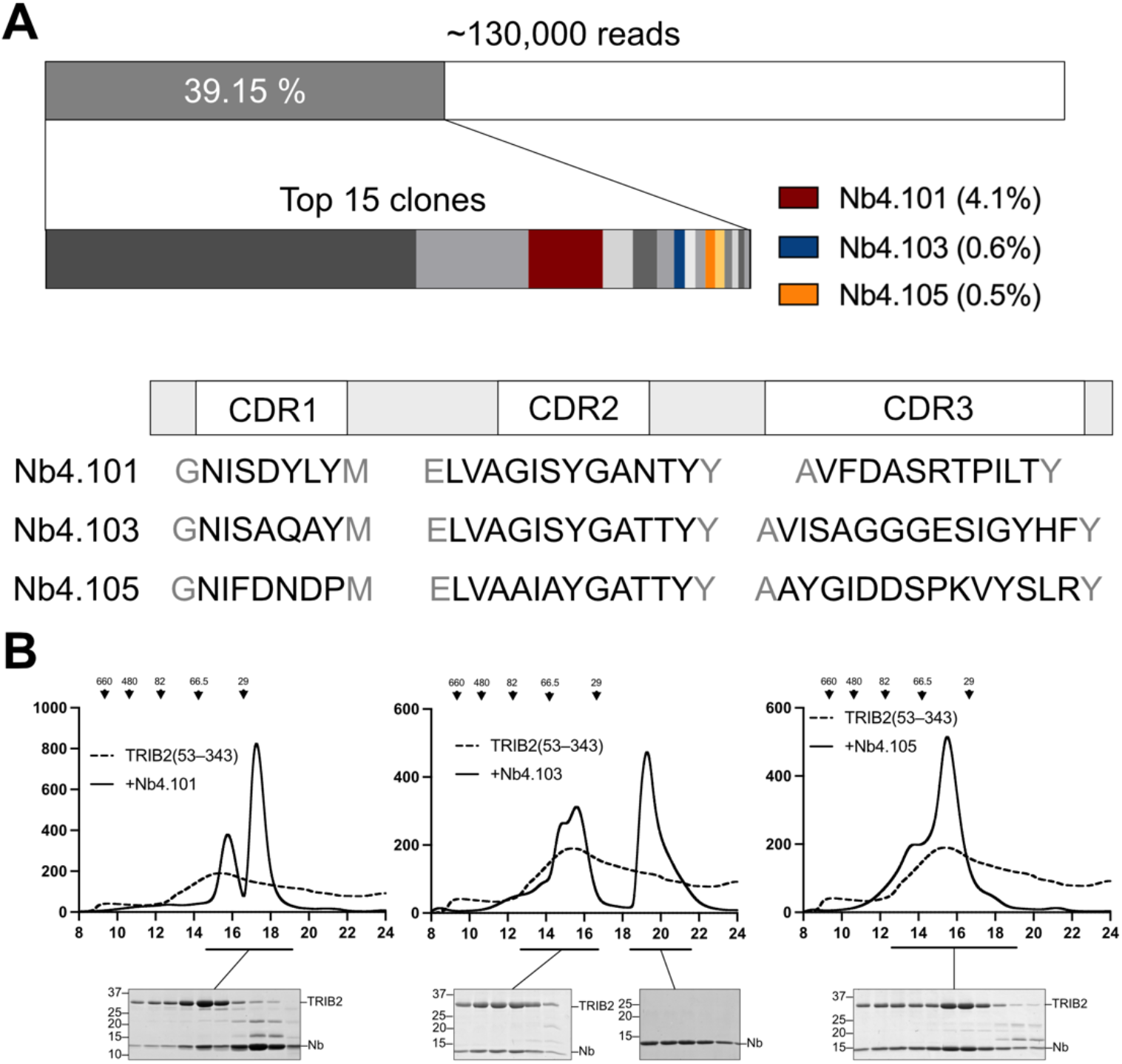
Identification of nanobodies that bind TRIB2. **(A)** Schematic representation of next-generation sequencing data. Fifteen clones made up almost 40 % of the reads. The proportions of the eventually selected clones as a percentage of total reads is indicated. **(B)** Size-exclusion chromatography of three selected nanobodies in combination with TRIB2(53–343). A trace representing TRIB2(53–343) alone is shown as a dotted line for comparison.

### Crystal structure of TRIB2-Nb4.103

Our previous attempts to determine the structure of the TRIB2 pseudokinase domain alone had been hampered by an inability to grow diffracting crystals and unstable protein. Having purified stable complexes of the three nanobodies in complex with TRIB2, we set about crystallisation screening. While small crystals of TRIB2-Nb4.101 grew, they were not readily improved. In contrast, initial crystals grew in multiple conditions for the TRIB2-Nb4.103 complex, especially when in complex with the core pseudokinase domain of TRIB2(53–313). Crystal optimisation allowed us to collect diffraction data extending to 2.7 Å resolution, and the structure of the TRIB2(53–313)-Nb4.103 complex was solved by molecular replacement (Figure 2A; Table 1). The structure contains a single TRIB2 and single nanobody in the asymmetric unit. Nb4.103 interacts with TRIB2 via the N-terminal lobe of the pseudokinase domain. In particular, the interface on TRIB2 is comprised of residues from the β-sheet comprised by β-strands 1–3. The nanobody side of the interface is largely contributed by residues from the CDR2 (complementarity-determining region 2) loop, with additional peripheral contributions from CDR1 and CDR3. Residues Thr56 (CDR2) and Tyr108 (CDR3) contribute to hydrogen bonds on either edge of the interface, while the core of the interaction is predominantly hydrophobic. The main contribution from CDR1 appears to be Tyr33 at the C-terminus of the variable region making hydrophobic interactions at the core of the interface (Figure 2B).

**Table 1:** Crystallographic data.

| Data Collection |  |
| --- | --- |
| TRIB2-Nb4.103 complex |  |
| Beamline | AS-MX2 |
| Wavelength (Å) | 0.9357 |
| Resolution (Å) | 2.70–45.7 |
| Space group | *P* 65 2 2 |
| Unit cell parameters<br>a, b, c (Å)<br>$\alpha$, $\beta$, $\gamma$ (°) | 75.160, 75.160, 273.770,<br>90, 90, 120 |
| R<sub>merge</sub> (outer shell) | 0.167 (1.306) |
| R<sub>pim</sub> (outer shell) | 0.070 (0.526) |
| Mean I/$\sigma$(I) (outer shell) | 10.2 (1.6) |
| Completeness (outer shell) | 100 (100) |
| Multiplicity (outer shell) | 6.3 (6.8) |
| Total No. of reflections | 85399 |
| No. of unique reflections | 13514 |
| CC(1/2) (outer shell) | 0.997 (0.502) |
| Wilson B factor (Å<sup>2</sup>) | 53.4 |
| Refinement Statistics |  |
| R<sub>cryst</sub> | 0.207 |
| R<sub>free</sub> | 0.260 |
| rmsd for bonds (Å) | 0.004 |
| rmsd for angles (deg) | 0.95 |
| rmsd for chiral volume (Å<sup>3</sup>) | 0.05 |
| No. of protein atoms | 2914 |
| No. of solvent atoms | 40 |
| Average B-factor overall (Å<sup>2</sup>) | 60.36 |
| Average main chain B-factor (Å<sup>2</sup>) | 58.2 |
| Average side chain and solvent B-factor (Å<sup>2</sup>) | 62.5 |
| Ramachandran plot statistics (%) |  |
| Favoured regions | 97 |
| Allowed regions | 3 |
| Outliers | 0 |

**Figure 2:**
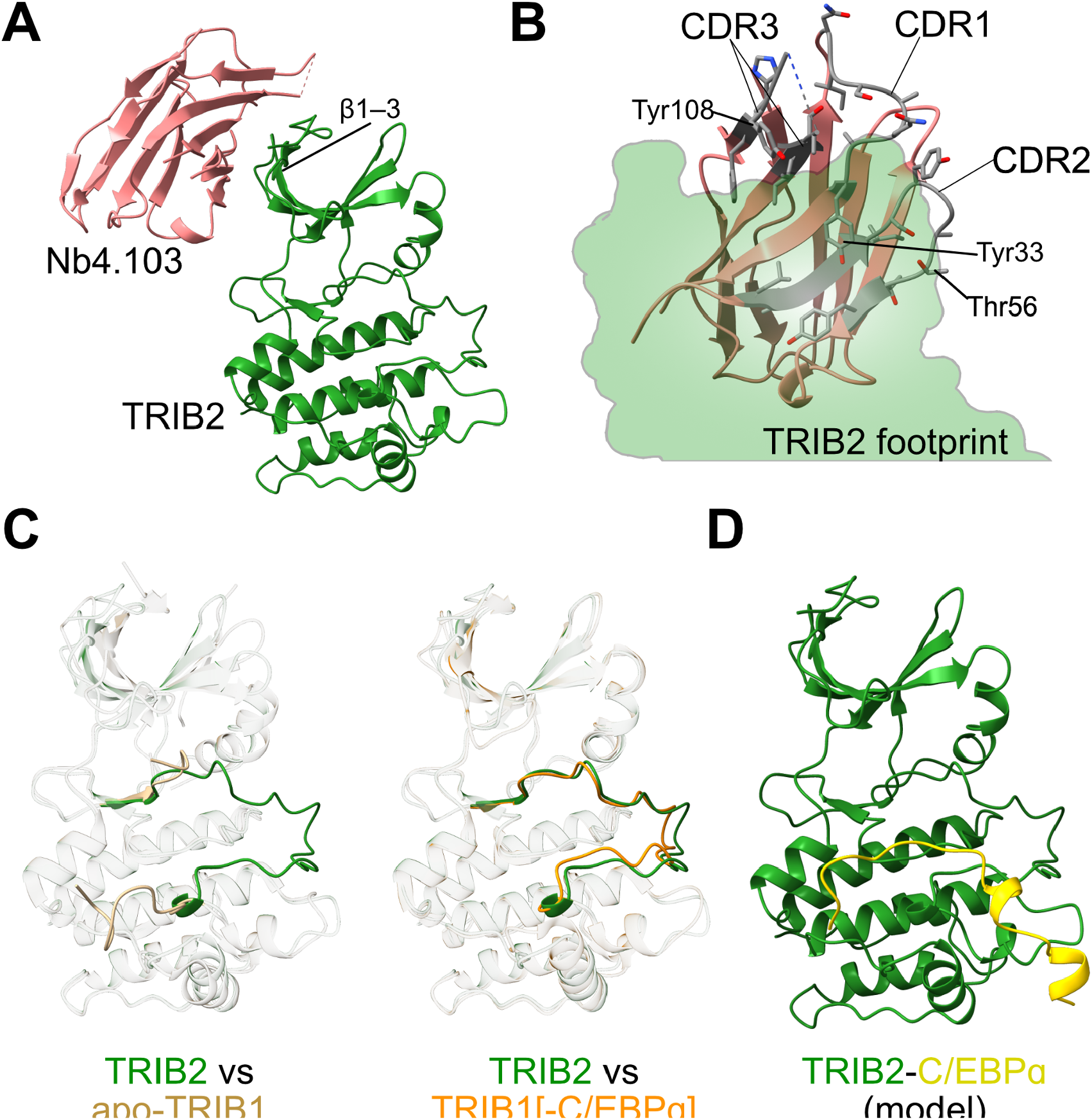
The TRIB2-Nb4.103 crystal structure. **(A)** Structure of the TRIB2-Nb4.103 complex. **(B)** View of the TRIB2-nanobody interface from the perspective of TRIB2. CDR regions are coloured grey, and residues mentioned in the main text are labelled. **(C)** Comparison of the TRIB2 structure with two different states of TRIB1. Left shows the superposition of TRIB2 (activation loop green) with the apo-state structure of TRIB1 without substrate (activation loop tan; PDB 5cem), whereas right shows the comparison with TRIB1 in the conformation bound to C/EBPa (activation loop orange; PDB 6dc0). **(D)** Model of the TRIB2 C/EBPa structure (green-yellow, respectively), created by superimposing PDB 6dc0 onto the TRIB2-Nb4.103 structure, and then showing only TRIB2 and C/EBPa.

The TRIB2-Nb4.103 structure was initially solved using a molecular replacement search model that consisted of the TRIB1 pseudokinase domain in the closed conformation, with the activation loop deleted (PDB code 5cem; (Murphy et al., 2015)). However, after model-building and refinement TRIB2 clearly adopts a more open conformation—whereby the activation loop is fully ordered, and does not block the region equivalent to the ATP-binding pocket of conventional kinases. This conformation is particularly reminiscent of the structure of TRIB1 bound to the C/EBPa substrate peptide, rather than the closed- or ‘apo’-form of TRIB1 (Figure 2C). Hence, the nanobody-stabilised structure appears to represent the activated state of TRIB2, where the TRIB2-C/EBPa complex can be modelled by simply superimposing the TRIB1-C/EBPa complex structure (Figure 2D). Beyond the C/EBPa binding-site, the TRIB2 structure closely aligns with the structure of TRIB1 (overall R.M.S.D of 0.835 Å over 222 residues). The bent aC-helix, and lack of a glycine-rich hairpin structure atop of the ATP-binding site that are characteristic of the TRIB1 pseudokinase domain are conserved in TRIB2, while the C-terminal lobe of the domain overlays well with both TRIB1 and conventional kinase domains. The structure also overlays well with the TRIB2 pseudokinase domain predicted by Alphafold2. This observation reinforces our observation that Alphafold2 tends to predict open activated conformations of TRIB1–3 with open active sites, as opposed to the closed conformation of TRIB1 that has been shown in two separate crystal forms.

### Comparison of TRIB1/2 pseudoactive sites

Previous studies of TRIB1 and TRIB2 have identified small-molecules that bind to the active site of each pseudokinase (or “pseudoactive” site) (Foulkes et al., 2018; Jamieson et al., 2018). Inhibitors that were originally designed to covalently target EGFR family tyrosine kinases have been shown to exhibit activity against TRIB2. Afatinib, in particular, covalently modifies Cys96 and Cys104 of TRIB2, which are within the aC-helix adjacent to the pseudoactive site (Figure 3A; (Foulkes et al., 2018)). Interestingly, there was limited overlap between TRIB2-targetting compounds and those that exhibited binding to TRIB1 (Foulkes et al., 2018; Jamieson et al., 2018). An experimental structure of TRIB2 now allows differences in inhibitor specificity to be rationalised.

**Figure 3:**
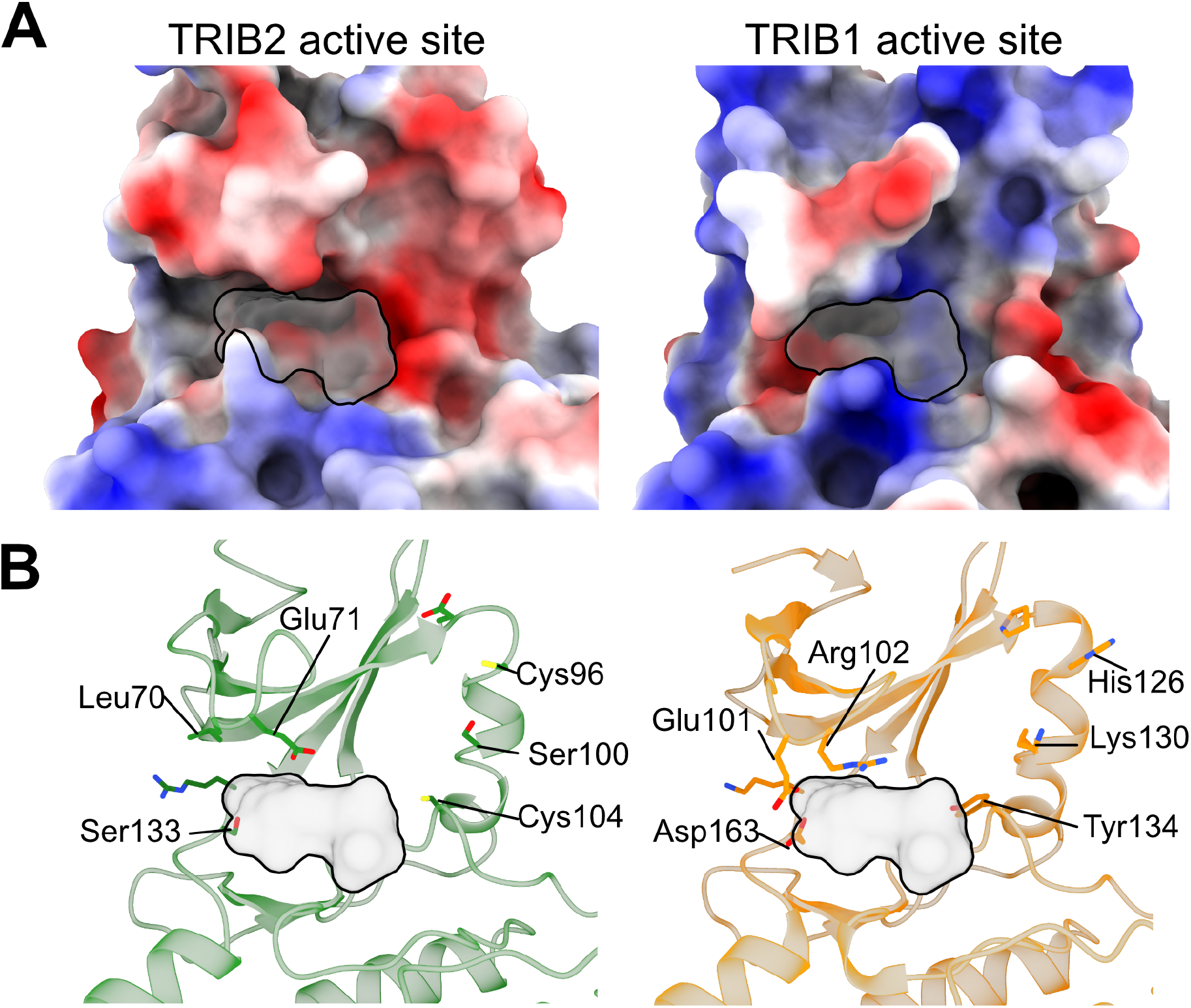
Character of the TRIB2 pseudoactive site relative to TRIB1. **(A)** Electrostatic surface of TRIB2 (left, this study) compared to TRIB1 (right, PDB code 6dc0) as calculated using default coulombic parameters in ChimeraX (red:electronegative–blue:electropositive). For reference, the position of an molecule in a conventional kinase is indicated in silhouette, based on superposition of with Protein Kinase A in complex with ATP and an inhibitory peptide (PDB 1atp; (Zheng et al., 1993)). **(B)** Cartoon representation of the same region shown in Panel A. Side chains of residues that differ between TRIB2 and TRIB1 are shown, and residues mentioned in the main text are labelled.

At a surface level, TRIB1 and TRIB2 exhibit surprisingly different electrostatic characteristics surrounding their pseudoactive sites (Figure 3A). TRIB2 offers a markedly more electronegative surface around the position where a conventional kinase would bind to ATP, mostly contributed by residues from the aC-helix (Figure 3A/B; Cys96–Ser100–Cys104). In contrast, these residues are comprised of Tyr134– Lys130–His126 in TRIB1, which contributes to a much more electropositive character. Adding to this charge disparity is Glu71 from the ‘Gly-rich’ loop above the active site of TRIB2, which occupies a similar position to the oppositely charged Arg102 of TRIB1. Perhaps most significantly is the location of Cys104 of TRIB2, located at the back of the active site pocket. Cys104 is the equivalent of Tyr134 in TRIB1, and it has been shown that Tyr-Cys mutation can destabilise TRIB1 more in line with the lower stability of TRIB2 (Jamieson et al., 2018). In the structure, Cys104 is solvent accessible—albeit somewhat occluded by Glu197 from within the activation loop, which is equivalent to the glycine within the DFG motif of conventional kinases. Thus, Cys104 offers a solvent accessible residue for modification by covalent inhibitors, which is also key to regulation of TRIB2.

Overall, these comparisons suggest that despite being the two most closely related Tribbles pseudokinases and having functional overlap, there could be significant scope for specific pharmacological selectivity between TRIB1 and TRIB2. The intimate relationship between regulatory residues in the pseudoactive site and substrate-binding of TRIB2, combined with cysteine reactivity at Cys104, could offer a route to inhibitors that specifically block TRIB2 substrate binding.

### Nanobodies selectively bind TRIB2 with nanomolar affinity

Because one downstream application of nanobodies is intracellular expression to modify protein function, we next sought to compare the affinity of different nanobodies for TRIB2, and test whether they could also recognise the pseudokinase domains of related Tribbles proteins TRIB1 and TRIB3. We first quantified the affinity of individual nanobodies for TRIB2 using isothermal titration calorimetry. As displayed in Figure 4A, each of the nanobodies exhibited tight binding characteristics—with dissociation constants of 57, 27 and 54 nM, for Nb4.101, Nb4.103, and Nb4.105, respectively. Interestingly, the nanobodies did show some variation in the calculated stoichiometry of the interaction. The stoichiometry of Nb4.101 and Nb4.103 titrated into TRIB2 was ∼0.5, indicating a ratio of two TRIB2 molecules per single nanobody per complex. In contrast, Nb4.105 showed a stoichiometry of N=1, indicating equal ratios of TRIB2 and nanobody per complex.

**Figure 4:**
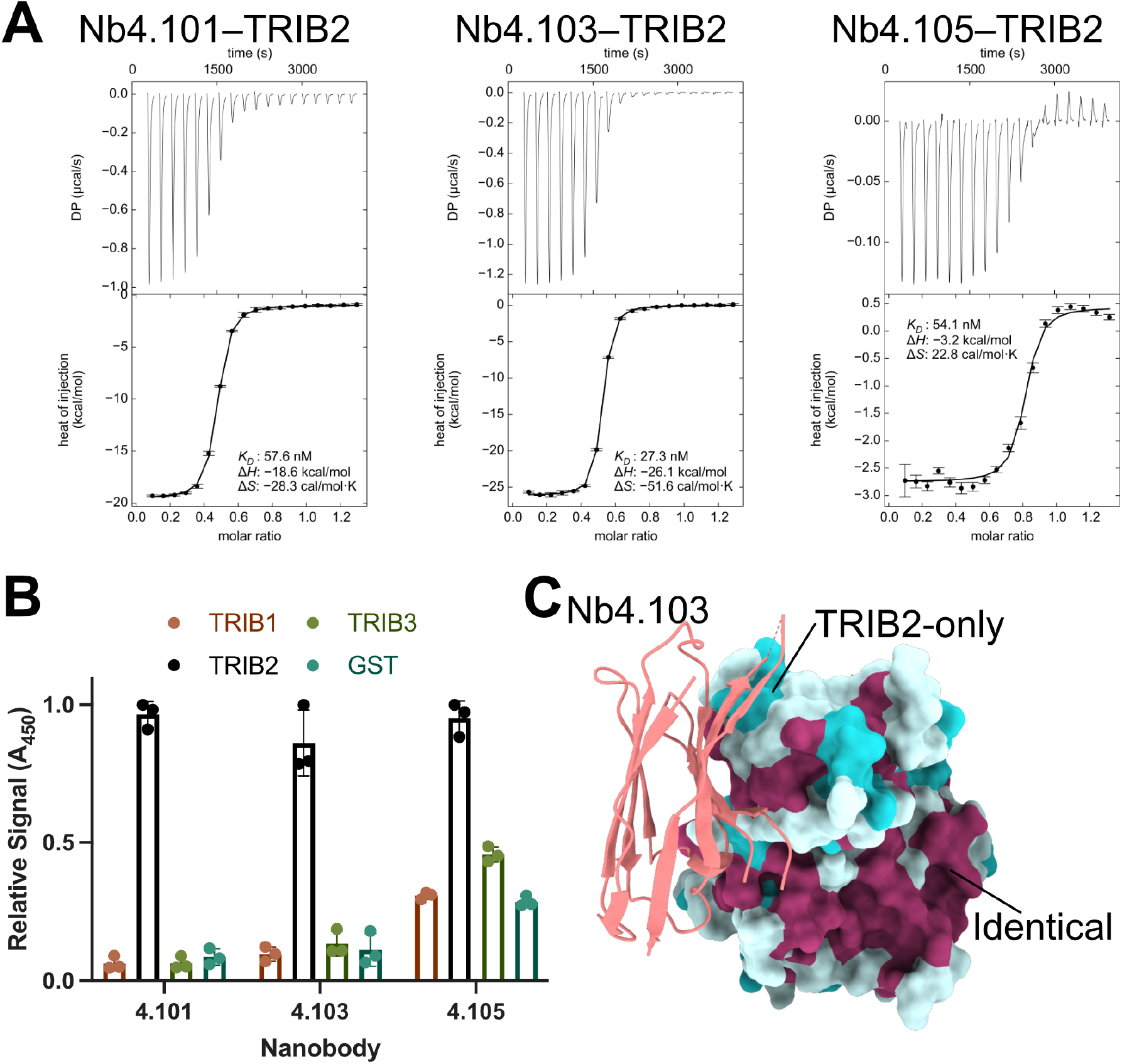
Binding properties of TRIB2 nanobodies. **(A)**Isothermal titration calorimetry of three respective nanobodies titrated into TRIB2(53–343). Thermodynamic properties of each titration are inset. **(B)** ELISA specificity assay of nanobodies against immobilised TRIB1, TRIB2, or TRIB3. Values are normalised to the maximum signal for each nanobody. **(C)** Conservation of the Nb4.103 surface of TRIB2. The TRIB2 surface is coloured according to a multiple alignment of the three human Tribbles proteins. Residues conserved in all three Tribbles are maroon, residues conserved in two of three proteins are light cyan, while residues that are different between the three Tribbles are dark cyan.

Within the pseudokinase domains, TRIB1 and TRIB2 are most closely related (71% identity based on pairwise comparison), whereas TRIB2 and TRIB3 are slightly more divergent (54% identity). TRIB1 and TRIB2 also share the conserved function of C/EBPa degradation. To test for nanobody cross-reactivity, we employed an enzyme-linked immunosorbent assay format, with Tribbles proteins (TRIB1–3) immobilised on the plate, and biotinylated forms of each nanobody that could be detected using HRP-conjugated streptavidin (Figure 4B). In these assays, Nb4.101 and Nb4.103 showed excellent selectivity for TRIB2 over TRIB1 or TRIB3, with mean signal against TRIB2 approximately 15–20-fold higher than signal against TRIB1 or TRIB3. Nb4.105 also showed preferential binding to TRIB2, albeit with lower selectivity (Figure 4B).

To rationalise the observed specificity of Nb4.103 for TRIB2, we compared the interface from the crystal structure with sequence conservation between the three Tribbles proteins. Mapping surface conservation shows that the epitope on TRIB2 is enriched in residues that are poorly conserved between the Tribbles orthologs (dark cyan), as opposed to residues that are identical between Tribbles (maroon). Altogether, these results suggest three different high-affinity nanobodies that selectively recognise natively folded TRIB2, over TRIB1 or TRIB3, and thus may form useful reagents for characterising TRIB2-specific functions in cells.

### Nanobody epitope analysis in solution

To ascertain if Nb4.101 and Nb4.105 bind to TRIB2 in similar modes to the crystal structure of TRIB2-Nb4.103, we employed an ELISA-based competition assay. In this format, unlabelled nanobodies were competed against biotin-labelled Nb4.103. Given all of the nanobodies had similar inherent affinity for TRIB2 (Figure 4A), it was expected that nanobodies with overlapping epitopes would successfully compete with 4.103, and as expected, unlabelled Nb4.103 was able to avidly compete with its labelled equivalent (Figure 5A). Nb4.101 was also equally able to compete with Nb4.103, indicating that it likely binds to an overlapping epitope on TRIB2. In contrast 4.105 was not able to displace Nb4.103, indicating a distinct epitope.

**Figure 5:**
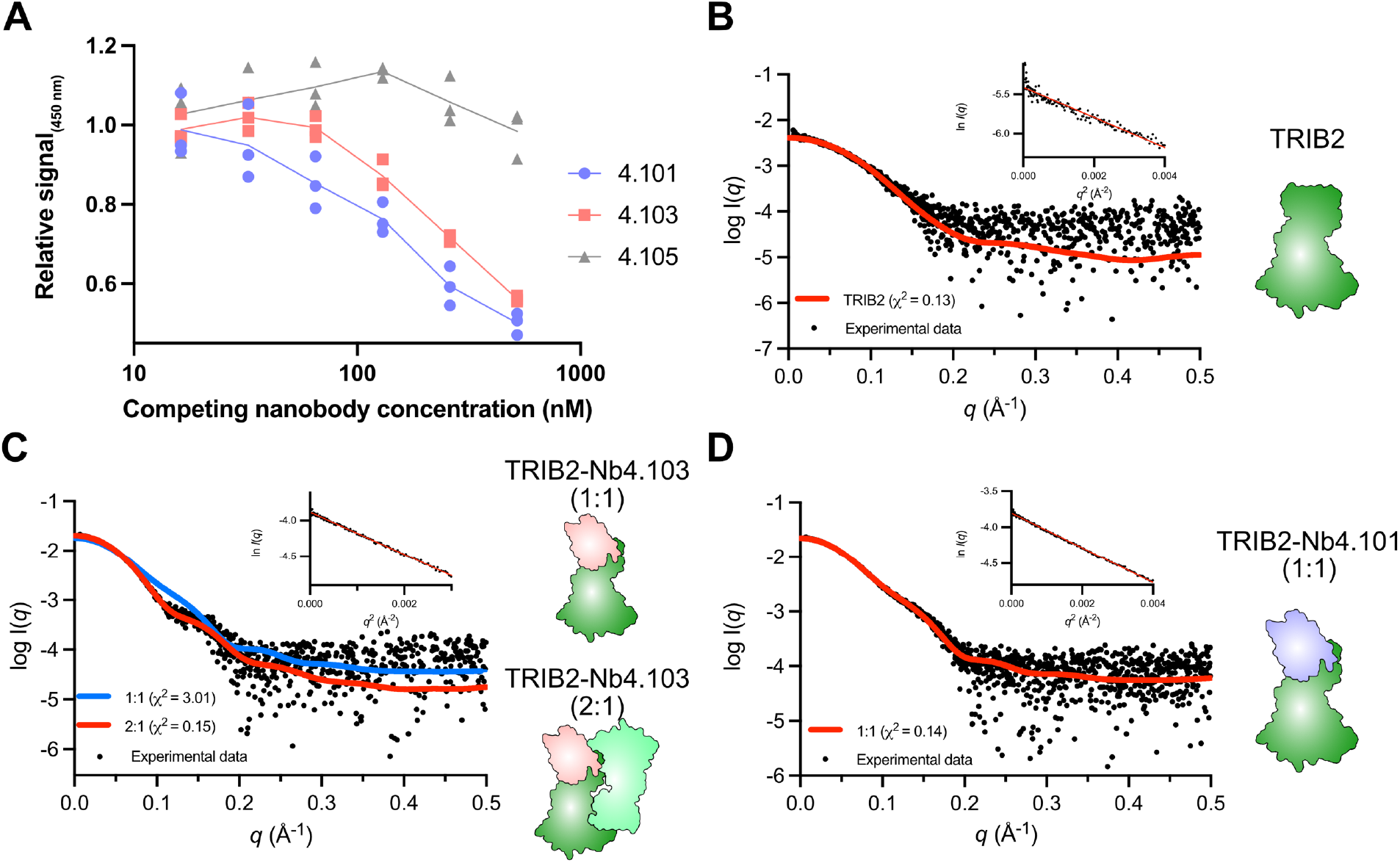
Epitope comparison of nanobodies. **(A)** Competition ELISA comparing ability of unlabelled nanobodies at indicated concentrations to displace biotinylated Nb4.103 from TRIB2. **(B)** Solution scattering profile of the TRIB2 structure as determined using SEC-SAXS (black circles). The experimental data is best fit to a monomeric model of TRIB2 alone (red line, χ^2^ = 0.13), shown in cartoon form. **C**) Solution scattering profile of the TRIB2-Nb4.103 structure as determined using SEC-SAXS. The experimental data is best fit to a model of a 2:1 complex of TRIB2-Nb4.103 (red line, χ^2^ = 0.15), rather than a 1:1 complex model (blueline, χ^2^ = 3.01). Models corresponding to each fitted complex, generated from the crystal structure are shown in cartoon form. **D**) Solution scattering profile of the TRIB2-Nb4.101 structure as determined using SEC-SAXS. The scattering data is best fit to a model of a 1:1 complex of TRIB2-Nb4.103 (red line, χ^2^ = 0.14), as shown in cartoon form. The Guinier plot is linear (inset, B-D), showing the sample is free from any measurable amounts of aggregation or inter-particle interference.

To test the binding modes of the TRIB2-Nb4.103 complex solution, we turned to size-exclusion chromatography coupled to small-angle X-ray scattering analysis (SEC-SAXS; Figure 5, Table 2, Supplementary Figure 1). As a control, scattering data from uncomplexed TRIB2 was an excellent fit for the monomeric apo-structure derived from the crystal structure (χ^2^=0.13), when the protein was analysed at the maximum concentration stable for the analyses (∼2.2 mg/mL/66 µM; Figure 5B, Table 2). Surprisingly, when loaded at concentrations from 1.7–7.16 mg/mL (37–159 µM) the TRIB2-Nb4.103 was not a good fit for the 1:1 complex observed in the crystal structure (χ^2^=3.01 using CRYSOL; Figure 5C). The *R*_g_ of the TRIB2-Nb4.103 complex scattering data suggested a mass of 70–79 kDa, more consistent with a 2:1 complex of TRIB2 with a single nanobody (Table 2). In analysing the interfaces between TRIB2 molecules in the crystal lattice using PISA, a face-to-face pseudokinase dimer that buried a surface area of 1513 Å^2^ was apparent (Krissinel and Henrick, 2007). Using the face-to-face TRIB2 dimer, a 2:1 TRIB2-Nb4.103 complex was an excellent fit for scattering data (χ^2^=0.15 using CRYSOL; Figure 5C), suggesting that Nb4.103 promotes a face-to face dimer of TRIB2. For Nb4.101, we did not see any indication of oligomerisation—the scattering data were an excellent fit (χ^2^=0.14 using CRYSOL; Figure 5D) for the 1:1 complex in the crystal structure when analysed up to a concentration of 6.2 mg/mL (137 µM) (Table 2). This indicates that the Nb4.101 and Nb4.103 have divergent properties in stabilising TRIB2 dimerisation, even though they bind to a similar epitope.

**Table 2:** SAXS data collection and analysis statistics.

| Data collection parameters |  |  |  |
| --- | --- | --- | --- |
| Instrument | Australian Synchrotron SAXS/WAXS beamline |  |  |
| Detector | PILATUS3-2M (Dectris) |  |  |
| Detector distance (m) | 3 |  |  |
| Wavelength (Å) | 1.0332 |  |  |
| Total $q$ range (Å<sup>-1</sup>)<sup>a</sup> | 0.005 – 0.6 | | |
| Maximum flux at sample | 8 x 10<sup>12</sup> photons per second at 12 keV |  |  |
| Exposure time | Continuous 1 second frame measurements |  |  |
| Sample configuration | SEC-SAXS with co-flow |  |  |
| Temperature (°C) | 12 |  |  |
| Analysis statistics | Trib2 | Trib2 Nb4.101 | Trib2 Nb4.103 |
| $I(0)$ (cm<sup>-1</sup>) (from Guinier analysis) | 0.004 ± 7.5e-05 | 0.022 ± 1.1e-04 | 0.017 ± 4.4e-05 |
| $R_g$ (Å) (from Guinier analysis)<sup>b</sup> | 23.49 ± 0.69 | 27.04 ± 0.22 | 30.12 ± 0.12 |
| $R_g$ (Å) (from $P(r)$ analysis) | 23.41 ± 0.41 | 27.21 ± 0.14 | 29.28 ± 0.14 |
| $D_{max}$ (Å) | 73 | 84 | 86 |
| Porod volume estimate (Å<sup>3</sup>) | 49911 | 61712 | 118176 |
| Molecular mass (MM) determination | Trib2 | Trib2 Nb4.101 | Trib2 Nb4.103 |
| MM (from Porod Volume, kDa) | 29.4 | 36.3 | 69.5 |
| MM (from Bayesian Inference, kDa) | 36.1 | 43.8 | 78.5 |
| Calculated monomer MM from sequence (kDa) | 29.9 | 42.9 | 42.9 |
| Estimated stoichiometry (Trib2:Nb4) | 1:0 | 1:1 | 2:1 |
| Software employed |  |  |  |
| Primary data reduction | ScatterBrain (Australian Synchrotron) |  |  |
| Data processing | PRIMUSQT (ATSAS) |  |  |
| Computation of model intensities | CRY SOL |  |  |

<sup>a</sup> $q$ is the magnitude of the scattering vector, which is related to the scattering angle ( $2\theta$ ) and the wavelength ( $\lambda$ ) as follows: $q = (4\pi/\lambda)\sin\theta$
<sup>b</sup> $R_g$ – Radius of gyration

The nature of the observed face-to-face TRIB2 dimer would block accessibility of the C/EBPa binding site on the front face of TRIB2. To test whether Nb4.103 may also influence C/EBPa binding, we measured their binding using isothermal titration calorimetry. The affinity of C/EBPa for TRIB2 was decreased approximately 3-fold in the presence of Nb4.103 (Figure 6A). In contrast Nb4.101 induced no such change in affinity. This suggests that the affinity of the face-to-face dimer is moderate, but enhanced in the presence of Nb4.103. The ITC experiments were executed with a TRIB2 concentration of ∼20 µM, whereas the observed TRIB2–C/EBPa dissociation constant is approximately 9 µM. In the concentration range of these experiments we hypothesise that substrate binding and TRIB2 dimerization exist in direct competition. In line with this hypothesis, we saw no difference in binding.

**Figure 6:**
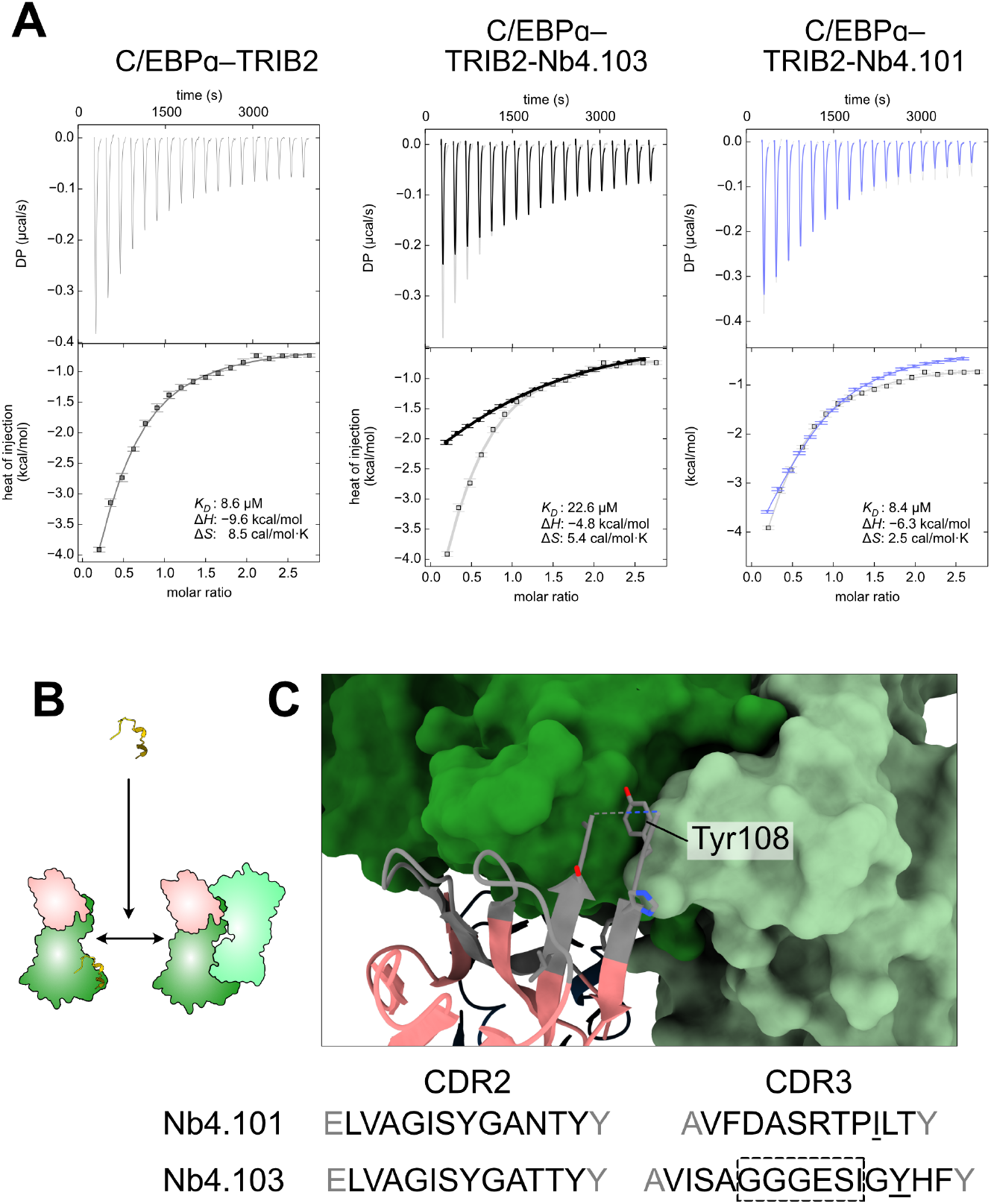
Effect of TRIB2 dimerisation on C/EBPa binding. **(A)** Isothermal titration calorimetry of MBP-C/EBPa titrated into TRIB2 alone (left), a TRIB2-Nb4.103 complex (centre) or a TRIB2-Nb4.101 complex (right). (B) Schematic representation of the interplay between C/EBPa binding and dimerization. Weak TRIB2 dimerization can be enhanced by Nb4.103 to an extent that reduces the apparent affinity of C/EBPa. (C) View of the TRIB2 dimer interface in relation to Nb4.103 CDR3. A comparison of CDR2 and CDR3 from Nb 4.101 and Nb4.103 is shown below, illustrating the relative conservation of CDR2 compared to CDR3. Tyr108 is labelled on the structure, and underlined in the Nb4.103 sequence, along with the equivalent isoleucine in Nb4.101.

Analysing the dimer interface in relation to TRIB2, provides a rationale for the observed oligomerisation behaviour. Nb4.101 and 4.103 share most of the same residues in CDR1 and CDR2, which is consistent with their shared overall epitope (Figure 5C). However, Nb4.103 has an extended CDR3 containing 13 residues, relative to 11 in Nb4.101. Although the glycine-rich repeat with CDR3 of Nb4.103 is not defined in the electron density, CDR3 sits near the interface between the two TRIB2 molecules, with Nb4.103 Tyr108 being wedged at the interface. The absence of a tyrosine residue and shortened CDR3 in Nb4.101 presumably contribute to its different oligomerisation behaviour to Nb4.103. These differences could facilitate distinct properties of the two nanobodies for modulating TRIB2 behaviour at high local concentrations.

## Discussion

Canonical protein kinases function with ATP bound in an active site between the N- and C-terminal lobes, correctly aligning catalytic residues for phosphoryl transfer. By virtue of eschewing catalytic function, divergence in these and surrounding sites are possible in pseudokinases, allowing them function in a variety of other ways (Jacobsen and Murphy, 2017; Mace and Murphy, 2021). For example, some pseudokinases with mutated catalytic residues can still bind nucleotides, and function by allosterically activating other kinases (HER3), behaving as signaling switches (MLKL), or acting as protein scaffolds (ULK4) (Garnish et al., 2021; Jura et al., 2009; Khamrui et al., 2020; Preuss et al., 2020; Shi et al., 2010). Alternatively, divergent pseudoactive sites can render pseudokinases incapable of binding ATP and/or magnesium, but still allow function across different modes of pseudokinase action (Mace and Murphy, 2021). Here we report the first experimentally-determined structure of the TRIB2 pseudokinase, and a trio of nanobodies that can be used for future studies of TRIB2 in biophysical and cellular experiments. The structure of TRIB2 differs from canonical kinase structural features in many of the same ways that have been observed for TRIB1. The aberrant glycine-rich loop and deformed aC-helix are each key TRIB-specific differences that contribute to an active site that is attenuated in binding ATP (Bailey et al., 2015b; Murphy et al., 2014, 2015). However, where the gross structural features surrounding the active site are similar, the physicochemical character of amino acids that form these structures is notably different. These specific differences suggest that small molecules that differentiate between the two proteins are a realistic possibility. Beyond small molecule inhibition, differences in pseudoactive site and activation loop residues could determine different dynamic tendencies of the two proteins, and determine different biological consequences.

Overexpression of either Trib1 or Trib2 from mouse bone marrow has been shown to drive AML (Dedhia et al., 2010). This could suggest that the two proteins are functionally redundant, given that they each function by binding to C/EBP family members and recruiting them for degradation by COP1. Based on the structure presented here, and measurement of affinity for C/EBPa, it appears that TRIB1 and TRIB2 can each bind to C/EBPa equally well. However, subsequent expression data has shown that TRIB1 and TRIB2 expression in humans has a reciprocal relationship during haematopoietic development—meaning high levels of TRIB1 coincide with low levels of TRIB2, and vice versa (Salomé et al., 2018a). The relevance of this reciprocal relationship is currently unclear, however different temporal patterns could mean that the two proteins are not simply playing the same role. One possibility is that while the two proteins share an ability to regulate C/EBPa, they each have distinct additional substrates that do not overlap, as illustrated by TRIB2’s role in regulation of P38 in cellular stress (Salomé et al., 2018b). The details of such a scenario likely requires targeted comparative proteomics to unravel. Alternatively, different functional effects could arise due to differences in dynamics of the two proteins, which alter turnover rates of common substrates such as C/EBPa, and thus transcriptional regulation.

Structures of TRIB1 have shown that the pseudokinase domain can also bind to its own C-terminal tail *in cis*, inhibiting itself from binding to COP1 (Murphy et al., 2015). Inherent to this mechanism is TRIB1 residue Tyrosine134, which sits at the base of the activation loop and at the back of the pseudoactive site pocket. It’s key role in switching between inactive and active states is illustrated in mutagenesis studies: where substituting the cysteine found at the equivalent position in TRIB2 (Cys104) destabilises TRIB1 (Jamieson et al., 2018). This suggests that TRIB2 may be more intrinsically more dynamic with its native cysteine residue in this position. From the structure it is not immediately clear why Nb4.103 promotes an activated state of TRIB2. However, its extensive interactions with β1–3 are likely to stabilise the entire N-terminal lobe of the kinase, which could have allosteric effects on the pseudoactive site and the C-terminal tail. While we were not able to solve the structure of TRIB2 including its C-terminal tail, it will be very interesting to determine if specifically modifying Cys104 could be the switch to regulate its interaction with COP1. Initial evidence suggests that this could be the case, with EGFR inhibitors that modify Cys96 and/or Cys104 of TRIB2 exhibiting off target activity and disruption of TRIB2 half-life (Foulkes et al., 2018). Developing more specific small molecules that regulate the conformational switch could be facilitated by the structural template that is provided through this work.

Reagents that can differentiate between the two most closely-related Tribbles proteins will be key to fully understanding the different functions of TRIB1 and TRIB2, and eventually to their targetability in disease. While nanobodies seem unlikely to be therapeutically viable option for Tribbles proteins, which are intracellular and expressed during differentiation—they could be extremely valuable as research reagents (Cheloha et al., 2020; Van et al., 2021). The three nanobodies we report here exhibit specificity for TRIB2 over both TRIB1 and TRIB3, and also differ in solution behaviour. The differences between Nb4.101 and Nb4.103 are particularly interesting given that Nb4.103 apparently promotes a more stable dimer of TRIB2 that blocks one of its key biological functions in binding to C/EBPa. It remains to be seen if a TRIB2 dimer has biological relevance, given that it appears to only occur at protein concentrations in the range of >10 µM. While these concentrations seem unlikely to occur in free solution in a cell, if TRIB2 binds to other proteins that oligomerise (potentially COP1) the local concentration of TRIB2 may increase to a point where the non-functional dimer may become more relevant.

The clear next step is to use TRIB2 nanobodies to probe biological function of the protein in a cellular setting. Nanobody affinity for TRIB2 is clearly sufficient for structure determination, but further validation is required to determine the types of application that may be possible. If the affinity and specificity are reflective of a preference for binding to TRIB2 in preference to other cellular proteins, we envisage that these nanobodies could offer uses in visualising TRIB2 by microscopy, flow cytometry, as affinity reagents, and/or modulating TRIB2 protein function through intracellular expression.

## Methods

### Protein expression and purification

All proteins (TRIB2 and nanobodies) were expressed in *E. coli* BL21(DE3) using Luria broth (Formedium) and nanobodies using auto-induction terrific broth (Formedium). Protein was first purified by immobilized metal-affinity chromatography (IMAC) using Ni^2+^ resin (Sigma), in a buffer consisting of 50 mM Tris pH 8.0, 300 mM NaCl, 10% sucrose, 10% glycerol, 10 mM imidazole. Protein was eluted using buffer supplemented with 300 mM imidazole. Tags were then cleaved while protein was dialysed overnight into 20 mM Hepes pH 7.6, 300 mM NaCl, 2 mM DTT —for His_6_-tagged tribbles protein cleavage was with 3C protease, or SUMO2 tagged nanobodies cleaved using SENP2 protease. Subsequent used to isolate TRIB2 were adopted from Jamieson et al. (2022). Namely the cleavage/dialysis step was performed in a temperature-controlled cabinet at 15°C, as TRIB2 constructs tended to precipitate overnight at 4°C. The following morning, proteins were passed back over Ni^2+^ resin to remove protease and uncleaved protein at room temperature, that had been equilibrated with dialysis buffer. Protein was then concentrated and further purified using size-exclusion chromatography (10/300 Superose-75 increase) in 20 mM Hepes pH 7.6, 300 mM NaCl, 2 mM DTT. Protein was aliquoted before flash-freezing in liquid nitrogen and storing at -80°C for later use.

For biotinylation, genes encoding nanobodies were sub-cloned with the N-terminal His_6_-SUMO fusion, into a pET-Duet vector adding a C-terminal avi-tag, and BirA enzyme co-expressed from a downstream multiple-cloning site (Middleton et al., 2021). These His_6_-SUMO-Nb-Avi fusions were then expressed using auto induction media supplemented with 200 µM biotin, and purified as above.

### Nanobody selection and characterisation

TRIB2 (53-343) was labelled with Alexa Fluor™ 647 and FITC before six initial screening rounds using MACS and FACS guided by the methods contained in McMahon et al (2018). The TRIB2 concentration was tapered during selection by interwoven magnetic and fluorescent cell sorting steps ((MACS AF647 1 µM; MACS FITC 500 nM; FACS AF647 250 nM; MACS FITC 100 nM; MACS AF647 30 nM; FACS AF647 1 nM). Illumina libraries were prepared for NGS from the 250, 30 and 1 nM-selected yeast libraries, and sequenced using 150 bp paired-end MiSeq reads. From these sequences, relative enrichment was used to prioritise nanobodies for characterisation. The top ten sequences were then synthesised (Geneart), cloned into a His_6_-SUMO2 vector and further characterised for on-column behaviour and using TRIB2 coelution.

### TRIB2-Nb4.103 crystallisation and data processing

For crystallization, TRIB2 (53-313) and nanobody 4.103 were pooled at a ratio of 1:1.5 (TRIB2: nanobody) based on nanodrop quantification following elution from nickel resin. The His_6_-tag from the Trib2 and the SUMO2 tag from the nanobody were then cleaved using 3C-protease and SENP2 protease and dialysed overnight at 15°C against 20 mM Hepes pH 7.6, 300 mM NaCl, 2 mM DTT. The following morning, the protein complex was washed over 2 mL HisSelect resin pre-equilibrated with dialysis buffer. The flowthrough was collected and concentrated to ∼15 mg.mL^-1^ before further purification by size-exclusion chromatography (10/300 S200 increase; 20 mM Hepes pH 7.6, 300mM NaCl, 2 mM DTT). Chromatography fractions were then analysed by SDS PAGE. While TRIB2 alone normally precipitates upon concentration after size-exclusion chromatography, whereas in complex with nanobodies it could concentrated to >10 mg.mL^-1^. Following concentration to ∼5 mg.mL^-1^, the TRIB2-Nb4.103 complex was aliquoted and snap frozen for storage at - 80°C.

Initial screening using TRIB2 (53–343)-NB4.103 complex yielded many crystals in numerous conditions from the Index HT, Peg/Ion HT, Clear strategy, and Morpheus screens. However, diffraction was limited or non-existent in most crystals. It was decided to rescreen using the shorter TRIB2 (53-313). Many crystals resulted in Index HT, Peg/Ion HT, Salt HT and Proplex screens. The crystals in PegIon HT B4 (0.2 M Magnesium nitrate hexahydrate, 20% w/v Polyethylene glycol 3,350) were most promising but diffraction was still poor. To further improve the crystals an additive screen was performed with PegIon B4 as the base condition. Crystals with 10 % sodium malonate additive were most promising, and used in a larger grid screen that yielded larger crystals. Crystals were cryoprotected in a mother liquor containing 25 % glycerol, and flash frozen.

Diffraction data was collected at the MX2 beamline at the Australian Synchrotron (Aragão et al., 2018). 600 frames were processed using XDS and Aimless (Kabsch, 2010; Winn et al., 2011). Molecular replacement was performed with the highest resolution TRIB1 structure (PDB 5cem; (Murphy et al., 2015)) and an exemplar nanobody generated from the same library (PDB 5vnv; (McMahon et al., 2018)). Cycles of manual rebuilding and refinement were performed using Phenix and COOT (Emsley and Cowtan, 2004; Liebschner et al., 2019). Analysis using UCLA server suggested anisotropy labelled as severe. Final structure refined with ellipsoidally-truncated data and a cycle of refinement optimisation using the PDB_REDO server (Joosten et al., 2014). Figures were prepared using UCSF ChimeraX (Pettersen et al., 2021). All software was installed and maintained using SBGrid (Morin et al., 2013).

### Isothermal titration calorimetry

ITC was performed essentially according to the methods in Jamieson et al. (2022).TRIB2 (53-343) and Nb4.101, Nb4.103 and Nb4.105 were defrosted and dialysed overnight against 20 mM Hepes pH 7.6, 300 mM NaCl, 0.5 mM tris(2-carboxyethyl)phosphine to ensure buffer matching. The TRIB2 and nanobody concentrations were adjusted to ∼20 µM and ∼120 µM respectively for each titration, and performed using a MicroCal VP-ITC. Titrations were analysed using NITPIC, SEDPHAT and GUSSI (Scheuermann and Brautigam, 2015; Zhao et al., 2015).

### Enzyme-linked immunosorbent assay

For ELISA analysis, a GST-control, GST-TRIB1(84–343), TRIB2(2–343) and MBP TRIB3(64–305) were purified and dialysed into 20 mM Hepes pH 7.6, 300 mM NaCl. Protein stocks were diluted to 65 nM, and 100 µL was added to each well of a MaxiSorp™ plate for immobilization. The plate was incubated overnight at 15°C with gentle shaking, then blocked for 120 min with 200 µL 20 mM Hepes pH 7.6, 300 mM NaCl, 0.05% Tween 20, 1% BSA at 15°C. The plate was washed three times with shaking for three minutes, 20 mM Hepes pH 7.6, 300 mM NaCl, 0.05% Tween 20. The biotinylated nanobodies were diluted to 65 nM and 100 µL was added to the plate. For the competition assay, the indicated dilution series of each nanobody was added with 65 nM of the biotinylated nanobody Nb4.103 simultaneously. Following incubation for 60 min shaking at room temperature, the plate was again washed three times. ExtrAvidin®−Peroxidase was then added and incubated with shaking covered for 60 min at room temperature. After washing, 90µL peroxidase solution-TMB and peroxidase substrates were mixed and added to each well and incubated at room temperature until the wells turned blue (around 15 min). Finally, 90 µL 0.1 M H_2_SO_4_ was added, and the plate was read at 450 nm. Analysis was started in Excel and final representations generated in GraphpadPrism.

### Small-angle X-ray scattering

SAXS data were collected at the Australian Synchrotron SAXS/WAXS beamline using an inline co-flow size-exclusion chromatography setup to minimize sample dilution and maximize signal-to-noise (Kirby et al., 2016; Ryan et al., 2018). Samples were analysed largely in-line with established methods (Meng et al., 2021; Oliver et al., 2021). Specifically, protein was injected (80 µL) onto an inline Superdex 200 5/150 Increase GL column (Cytiva), equilibrated with size exclusion buffer (50 mM Tris-HCl, pH 8, 300 mM NaCl, 5% glycerol) at 12 °C, at a flow rate of 0.2 mL min^−1^. 2D intensity plots were radially averaged, normalized against sample transmission, and background-subtracted using the Scatterbrain software package (Australian Synchrotron). The ATSAS software package was used to perform the Guinier analysis (PrimusQT; (Konarev et al., 2003)), to calculate the pairwise distribution function *P*(*r*) and the maximum interparticle dimension (*D*_max_), and to evaluate the solution scattering against the crystal structure solved in this study (CRYSOL; (Svergun et al., 1995)). All models were generated using ChimeraX. All data collection and processing statistics are summarised in Table 2.

## Acknowledgements

We are especially grateful for the yeast nanobody library made available by Andrew Kruse (Harvard University) and Aashish Manglik (University of California, San Francisco). We thank Patrick Eyers (University of Liverpool) for original gift of TRIB2 DNA template and TRIB3 expression plasmids, and Karen Keeshan for comments on the manuscript. The BirA co-expression plasmid for biotinylation was a kind gift from Adam Middleton (University of Otago). We acknowledge the facilities, and scientific and technical assistance from flow cytometry staff at Otago Micro and Nanoscale Imaging (OMNI), at the University of Otago. We are grateful to the staff of the Australian Synchrotron SAXS/WAXS beamline for their assistance with data collection. Diffraction data was collected at the MX2 beamline at the Australian Synchrotron, part of ANSTO, and made use of the ACRF detector and thank the New Zealand synchrotron group for facilitating access to the MX beamlines. This work was supported by the Marsden Fund Council from New Zealand Government funding, managed by Royal Society Te Apārangi. JMM gratefully acknowledges the support of the National Health and Medical Research Council of Australia (fellowship, 1172929; IRIISS, 9000719) and the Victorian Government Operational Infrastructure Support scheme.

---

**Supplementary Figure 1.**
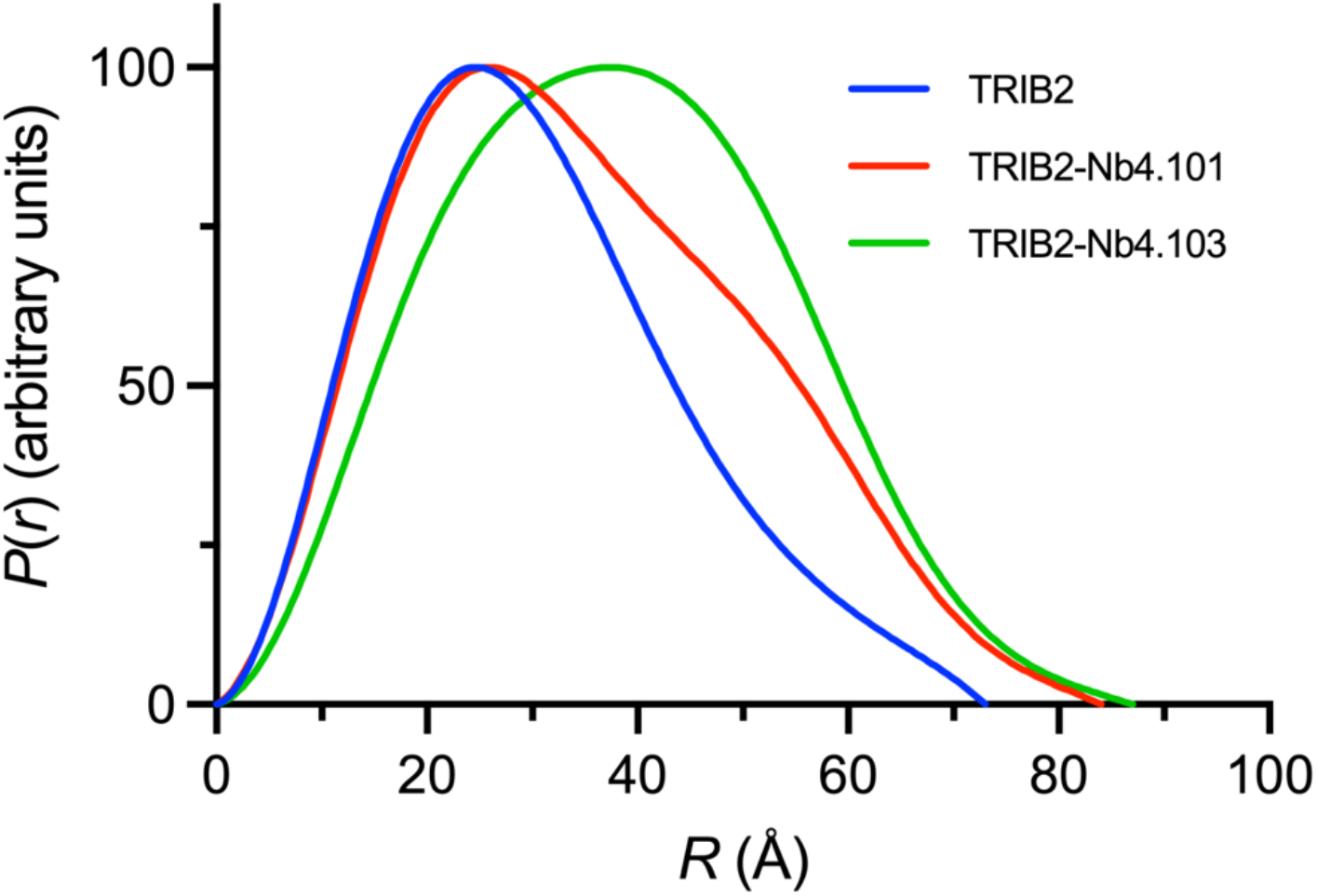
Interatomic distance distributions of TRIB2(53–343) and nanobody complexes.

## Supplementary Data

*Full nanobody sequences (CDR regions underlined)*

***4*.*101***

QVQLQESGGGLVQAGGSLRLSCAASG<u>NISDYLY</u>MGWYRQAPGKERE<u>LVAGISYG ANTY</u>YADSVKGRFTISRDNAKNTVYLQMNSLKPEDTAVYYCA<u>VFDASRTPILT</u>YWG QGTQVTVSS

***4*.*103***

QVQLQESGGGLVQAGGSLRLSCAASG<u>NISAQAY</u>MGWYRQAPGKERE<u>LVAGISYG ATTY</u>YADSVKGRFTISRDNAKNTVYLQMNSLKPEDTAVYYCA<u>VISAGGGESIGYHF</u> YWGQGTQVTVSS

***4*.*105***

QVQLQESGGGLVQAGGSLRLSCAASG<u>NIFDNDP</u>MGWYRQAPGKERELV<u>AAIAYG ATTY</u>YADSVKGRFTISRDNAKNTVYLQMNSLKPEDTAVYYCA<u>AYGIDDSPKVYSLR</u> YWGQGTQVTVSS

